# Posterior but not frontal neural signatures of subjective visibility in report-independent EEG decoding

**DOI:** 10.64898/2026.04.20.719596

**Authors:** Sabine Gnodde, Vlada Aslanov, Jolien C. Francken, Abigail Hogan, Umberto Olcese, Timo Stein

## Abstract

A central question in consciousness research is whether perceptual awareness arises predominantly from activity in posterior sensory regions or from later frontal processes associated with access and report. Empirical tests of these alternatives are challenging because most paradigms rely on explicit perceptual reports, introducing motor and decisional confounds. Here, we used an electroencephalography design combining two independent tasks to isolate neural signatures of subjective visibility while minimizing report-related activity. Participants viewed lateralized gratings in a backward-masking task and rated their subjective visibility, while in a separate no-report task they viewed unmasked gratings without reporting. Classifiers were trained on stimulus location in the no-report task and tested on masked trials sorted by subjective visibility in the masking task. This cross-decoding approach isolates stimulus-specific neural representations that generalize across tasks and are therefore independent of reporting and decision processes. Decoding revealed a reliable difference between subjectively visible and invisible stimuli in an early time window from 130 to 170 milliseconds, driven by posterior sensors over occipital, temporal, and parietal regions. No corresponding effects were observed over frontal sensors. These results indicate that subjective awareness is associated with early, posterior neural representations, whereas frontal activity observed in report-based paradigms likely reflects post-perceptual processing rather than awareness per se.

## Introduction

Identifying which neural circuits support conscious perception remains a central challenge in neuroscience. Research on the neural correlates of consciousness (NCCs) aims to identify the minimal neural activity sufficient for conscious contents to arise (Crick & Koch, 1990). A common empirical strategy is to compare brain activity between conditions in which the same stimulus is consciously perceived versus not perceived, using paradigms that manipulate visibility while holding stimulus input constant. Such paradigms include binocular rivalry (Blake et al., 2014; Li et al., 2017), attentional blink (Sergent et al., 2005; Slagter et al., 2017), and visual masking (Dehaene et al., 2001; Kouider & Dehaene, 2007). Findings from these paradigms have motivated competing theories of consciousness that differ in their claims about where and when conscious contents emerge: either through late, global access mechanisms (Dehaene & Changeux, 2011; Lau & Rosenthal, 2011; Mashour et al., 2020) or earlier, recurrent processing within posterior cortex (Lamme, 2006; Tononi, 2008, 2012, 2015). Recent adversarial collaborations have sought to adjudicate between these competing frameworks empirically (COGITATE Consortium et al., 2023; Ferrante et al., 2025), yet their conclusions remain contested.

A major reason for this impasse is that many neural markers traditionally associated with consciousness may instead reflect post-perceptual processes. In report-based paradigms, participants must decide, respond, and act on their perceptual experiences. These processes engage attention, working memory, decision-making, and motor preparation, which recruit fronto-parietal circuits that overlap with proposed NCCs (Aru et al., 2012; Naccache et al., 2015). As a result, neural activity correlated with conscious reports may partly reflect access and reporting rather than visual awareness.

To address this problem, no-report paradigms have been developed in which perceptual awareness is inferred from behavioral or physiological markers without requiring explicit reports (Cohen et al., 2020, 2024; Tsuchiya et al., 2015). These approaches have reduced or eliminated many frontal and late components, suggesting a more posterior locus of perceptual processing. However, no-report paradigms have their own limitations: it is often unclear whether stimuli were consciously perceived on a given trial (Naccache et al., 2015), participants may still internally monitor their experience (“bored monkey problem”; Block, 2019), and some studies continue to report residual frontal or late activity (Cohen et al., 2020; Hatamimajoumerd et al., 2022). As a result, it remains debated whether prefrontal involvement reflects awareness or post-perceptual processes driven by task structure.

Here, we introduce a hybrid approach designed to combine the strengths of report and no-report paradigms while minimizing their respective weaknesses. Participants performed a backward masking task in which lateralized gratings were briefly presented and followed by a backward mask, and subjective visibility was assessed using the Perceptual Awareness Scale (PAS; Ramsøy & Overgaard, 2004). In a separate, independent no-report task, the same stimuli were presented clearly and without masking, and no perceptual reports were required.

We used multivariate pattern analysis (MVPA) to decode stimulus location from electroencephalography signals, training classifiers on the no-report task and testing them on the masking task. This cross-decoding strategy isolates stimulus-specific neural representations that generalize across tasks, thereby minimizing contributions from motor responses, decision processes, and report-related activity while still allowing trials to be labeled by subjective visibility. By decoding stimulus location as a proxy for content-specific perceptual representations, we could compare neural responses between subjectively visible and invisible stimuli that were physically identical. This approach allows us to directly test whether conscious perception depends on late, fronto-parietal global access mechanisms or earlier, posterior recurrent processing, as reflected in the timing and topography of decoding.

In our study, we identified a reliable 130 to 170 millisecond time window in which stimulus location could be decoded more accurately for subjectively visible than invisible trials, with contributions from posterior electrodes and no evidence for prefrontal involvement. These results indicate that subjective visibility is associated with early, posterior neural representations that generalize across report contexts, supporting posterior and recurrent accounts of conscious perception while challenging theories that locate the NCC in prefrontal cortex.

## Methods

### Participants

The experimental protocol was approved by the University of Amsterdam Ethics Committee (approval number 2019-BC-15504). All participants provided written informed consent and were compensated for their time. Participants reported normal or corrected-to-normal vision and were naïve to the research question. Initially, 108 participants were recruited. Fourteen were excluded due to technical issues, incomplete data, or discontinuation (e.g., EEG setup difficulties). Three additional participants were excluded due to a coding error that led to unbalanced conditions. This resulted in a final sample of 91 participants (78% female, mean age = 21.25 years, SD = 2.51, age range 17–29 years). All remaining participants were included in analyses, except for the main task k-folding procedure, where an additional 8 were excluded due to insufficient trial numbers, resulting in N = 83. This sample size exceeds those of previous related studies (Cohen et al., 2020; Krisst & Luck, 2025; Motlagh et al., 2024) and is therefore considered sufficient.

### Stimuli and presentation

Target stimuli were circular black-and-white Gabor patches (6° diameter, 0.89 cycles/°, phase = 0.25 with Gaussian smoothing, 4.5 cm, 40% contrast). Each target was presented in either a vertical or horizontal orientation and appeared in the center of the left or right half of the screen (8° to the left or right of fixation, 0° vertically). Masks were created by superimposing horizontal and vertical Gabor patches of the same size as the target (50% contrast) and were shown in the same locations as the targets on both sides of the screen. All stimuli were displayed on a 24-inch LCD screen (1920x1080 pixel resolution, 120-Hz refresh rate) against a grey background (RGB: R127.5, G127.5, B127.5). Stimulus presentation was controlled using MATLAB and Psychtoolbox (Brainard, 1997). Participants viewed the screen from approximately 60 cm in a dim, sound-attenuated room, with their heads stabilized using a chin rest.

### Experimental procedure

The study consisted of two separate experiments: (1) a main masking task requiring explicit perceptual reports, and (2) an independent no-report task involving passive viewing without reports. Prior to the experiment, participants received detailed written and verbal instructions, were asked to sit still, fixate on a central cross, avoid blinking, and keep both eyes open. They were informed that the target would appear on either the left or right side of fixation and that its visibility would vary due to masking. Responses were self-paced, with no time limit, and the next trial was initiated by a keypress. Stimulus timings were synchronized to the screen’s refresh rate to ensure precision. Throughout both tasks, whole-brain EEG data were recorded for multivariate analyses.

#### Main masking task

The main masking task was designed to obtain visibility conditions that were defined both subjectively (i.e., based on subject’s visibility report) and objectively (i.e., based on subject’s stimulus localization accuracy), using a backward masking paradigm with varying stimulus onset asynchronies (SOAs). Each trial began with a grey screen and a white fixation cross for 1000 ms, followed by a jittered interval of 300–600 ms. A target stimulus (8 or 16 ms) then appeared on either the left or right side, followed by a mask presented for 200 ms. SOAs between target and mask varied across four conditions: 8 ms, 50 ms, 150 ms, and a mask-only control condition in which no target was shown (Fig. 1A). Each SOA condition consisted of 288 trials (20% of total), except the 50 ms SOA condition, which included 576 trials (40%) to allow later subdivision based on PAS ratings. In total, the experiment contained 1440 trials, divided into 16 blocks with breaks.

**Figure 1.**
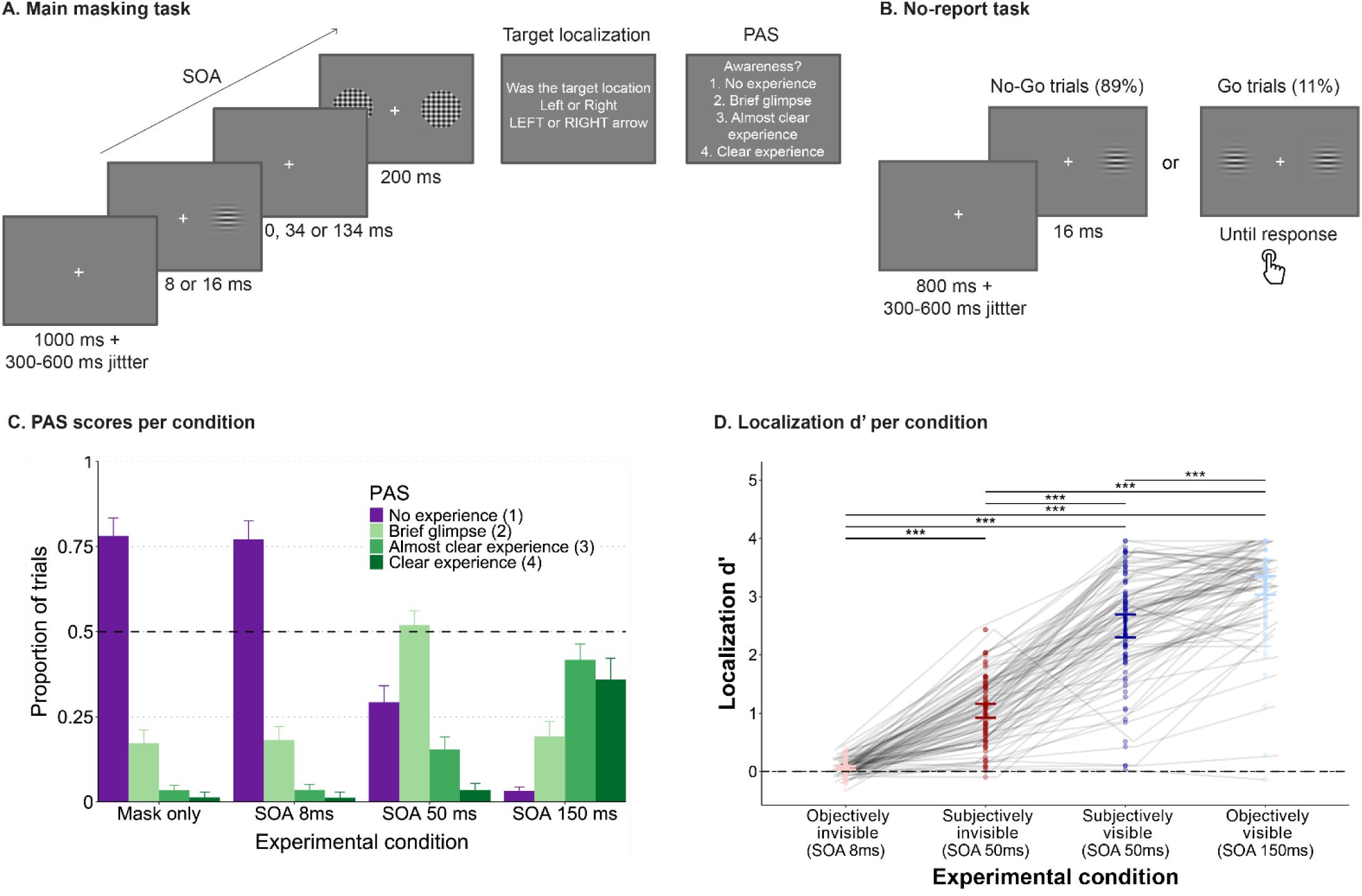
Behavioral paradigm and results. **(A)** Main masking task. A grating stimulus was presented for 8 or 16 ms and followed by a mask after a variable stimulus onset asynchrony (SOA; 8, 50 or 150 ms). Participants reported the stimulus location and subsequently rated their subjective awareness using the perceptual awareness scale (PAS). **(B)** No-report task (i.e., stimulus localizer). In this passive viewing paradigm, single targets were presented on the left or right during No-Go trials, requiring no response. On 11% of trials (Go trials), two targets were presented and participants responded with a keypress. Data from this task were used to train classifiers for cross-decoding analyses. **(C)** PAS ratings. Group-averaged proportion of trials per stimulus condition (mask only, 8 ms SOA, 50 ms SOA, and 150 ms SOA) as a function of PAS rating: no experience (PAS = 1), brief glimpse (PAS = 2), almost clear experience (PAS = 3), and clear experience (PAS = 4). Error bars indicate SEM. **(D)** Localization accuracy. Individual and group-averaged localization sensitivity (d′) for discriminating stimulus location (left vs. right) across conditions: 8 ms SOA, 50 ms SOA without awareness (PAS = 1; “subjectively invisible”), 50 ms SOA with awareness (PAS = 2–4; “subjectively visible”), and 150 ms SOA.

After each trial, participants were first asked to indicate the location of the target (left/right) using the right-hand arrow keys. If they did not perceive a stimulus, they were instructed to guess. Next, they used their left hand to rate subjective experience on the PAS: (1) no experience, (2) brief glimpse, (3) somewhat clear experience, or (4) a completely clear experience (Ramsøy & Overgaard, 2004).

#### No-report task

The no-report task provided EEG data for classifier training in the absence of explicit perceptual reports. The same target stimuli were presented, but without masking and without requiring responses regarding stimulus location or subjective experience.

Each trial began with a grey screen and white fixation cross for 800 ms, followed by a jittered interval of 300–600 ms. In 89% of trials, a single target appeared on the left or right (no-response trials), and participants were instructed to passively view the stimulus. To keep participants engaged, the remaining 11% of trials were response-trials in which two target stimuli appeared simultaneously on both sides of fixation. Participants were instructed to respond to these response-trials with a keypress. These response-trials were excluded from all analyses to avoid motor- or attention-related confounds (Fig. 1B).

### EEG acquisition and preprocessing

EEG was recorded using a 64-channel BioSemi ActiveTwo system (www.biosemi.com) at a sampling rate of 1024Hz. Two earlobe electrodes served as reference channels and four electro-oculographic (EOG) electrodes were used for monitoring eye movements and blinks.

Preprocessing was performed in MATLAB (R2023b, MathWorks) using the FieldTrip toolbox (Oostenveld et al., 2011) and EEGLAB (Delorme & Makeig, 2004). Raw .bdf files were converted to .set format, allowing merging of interrupted sessions and trigger correction. The EEG signal was re-referenced to the earlobe average, downsampled to 128 Hz to reduce computational load, epoched from –250 to 1000 ms relative to stimulus onset, and baseline-corrected using the –250 to 0 ms interval.

### Data Analysis

#### Behavioural data

Behavioural analyses were conducted in RStudio (Version 4.4.0; The R Foundation for Statistical Computing) while all other analyses were performed in MATLAB (Version R2023b, MathWorks). For each main task condition (mask only, 8 ms SOA, 50 ms SOA, and 150 ms SOA), the proportion of trials corresponding to each PAS response (scores 1–4) was calculated. The 50 ms SOA trials were split into two categories based on subjective awareness: subjectively visible trials (PAS 2–4) and subjectively invisible trials (PAS 1). This enabled classification of trials based on subjective awareness on a trial-by-trial basis, while the physical stimulus remained the same.

Objective accuracy was assessed using the sensitivity index d-prime (d’), computed for each visibility condition (8 ms SOA, 50 ms subjectively visible, 50 ms subjectively invisible, 150 ms SOA) based on localization hit and false alarm rates. The objectively invisible and objectively visible conditions served as reference conditions for assessing neural responses under minimal and maximal perceptual awareness, respectively. Thus, two experimental conditions and two reference conditions were defined:

1. Subjectively invisible: Target ON 16 ms; Target OFF 34 ms; SOA 50 ms; PAS= 1
2. Subjectively visible: Target ON 16 ms; Target OFF 34 ms; SOA 50 ms; PAS= 2–4
3. Objectively invisible (reference): Target ON 8 ms; Target OFF 0 ms; SOA 8 ms
4. Objectively visible (reference): Target ON 16 ms; Target OFF 134 ms; SOA 150 ms

#### Multivariate Pattern Analysis (MVPA)

Stimulus location (left vs. right) was decoded from EEG activity using the Amsterdam Decoding and Modeling (ADAM) toolbox (Fahrenfort et al., 2018) in MATLAB, which supports both single-subject and group-level MVPA. At the single-subject level, a backward decoding model with Linear Discriminant Analysis (LDA) was used to classify stimulus location based on EEG patterns. To ensure balanced training, both within-class balancing (equal number of left/right trials) and between-class balancing (equal representation of experimental condition) were applied. Decoding performance was assessed using d’. Two decoding strategies were used (Table 1):

1. **Main task K-Folding:** 10-fold cross-validation with 90% training and 10% testing, repeated for all main task conditions and used for comparison with cross-decoding analyses.
2. **Cross-decoding:** Classifier trained on the no-report task and tested on main task conditions, isolating neural activity related to perceptual processing independent of post-perceptual processes, as training data contained no reports.

**Table 1.** Decoding strategies for conscious content used in multivariate pattern analysis (MVPA)

| Decoding strategy | Training data | Testing data | Advantages | Disadvantages |
| --- | --- | --- | --- | --- |
| **Main task K-Folding** | Main task (incl. subjective visibility) | Main task (incl. subjective visibility) | Reflects subjective visibility | Post-perceptual processing effects |
| **Cross-decoding** | No-report task | Main task (including subjective visibility) | Reflects subjective visibility independent of post-perceptual processing effects | - |

Initial decoding used all 64 EEG channels, taking advantage of MVPA’s ability to detect patterns without a priori electrode selection. The classifier was trained on data from −250 to 1000 ms relative to stimulus onset for every sample in the epoch. To further examine spatial contributions, region-by-region cross-decoding was performed using predefined electrode pools, allowing investigation of the involvement of different cortical regions.

To localize neural sources contributing to decoding, classifier weight vectors were transformed into activation patterns using the data covariance matrix following Haufe et al., 2014. This approach multiplies the classifier weights by the data covariance matrix to produce interpretable activation patterns that can be directly interpreted as neural sources. These were averaged within four group-defined time windows (80–190 ms, 190–260 ms, 260–430 ms, 430–900 ms), derived from group-level decoding peaks, enabling examination of both temporal and spatial dynamics associated with conscious perception.

For both the main task K-folding and the cross-decoding analyses, decoding performance was quantified using d′. Although the AUC metric was also computed (Suppl. Fig. 1), this did not reach significance, likely due to the brief and transient nature of the effect. Because d′ directly captures the separation between signal and noise distributions, it is better suited for detecting subtle effects under low signal-to-noise conditions. Importantly, the significant d′ cluster was robust across 20 permutation runs, confirming it was not driven by chance.

#### Temporal generalization

Temporal generalization was used to assess whether neural patterns generalize across time, testing whether a classifier trained at one time point can distinguish left vs. right stimulus locations at other time points (King & Dehaene, 2014). At the group level, averages of decoding performance (d’) were computed to characterize results and compare main task K-folding and cross-decoding results. While diagonal decoding assessed classifier accuracy at each individual time point, temporal generalization matrices (TGMs) captured the stability of neural representations over time. D’ was tested against chance and corrected for multiple comparisons via cluster-based permutation tests.

#### Statistical tests

Statistical significance was assessed using two-tailed t-tests with cluster-based permutation correction for multiple comparisons. For decoding, classifier performance was compared to chance (50%) to determine significant decoding of stimulus information. Condition differences were assessed using subject-level t-tests on d′ values at each time point, with statistical significance defined as p < 0.05. To address the multiple-comparisons across time, we employed cluster-based permutation testing, in which clusters were identified as sequences of temporally contiguous samples exceeding the significance threshold.

To complement the region-by-region analyses, we performed Bayesian paired-sample t-tests at each time point on subject-wise decoding accuracy differences (visible - invisible). Bayes factors (BF₁₀) quantified evidence for a directional alternative (H_+_: δ > 0) versus the null (H₀: δ = 0), using a half-Cauchy prior on the standardized effect size (scale r = 0.707; Rouder et al., 2009). BF were computed independently at each time point, uncorrected for multiple comparisons and summarized within the predefined 75–180 ms window using the median BF₁₀ following Jeffreys–Wagenmakers conventions. Bayesian analyses were conducted specifically for the region-by-region comparisons to quantify evidence for the absence of a visibility effect, particularly in frontal electrodes, where frequentist analyses yielded non-significant results. This approach allowed us to distinguish between lack of sensitivity and evidence supporting the null hypothesis.

## Results

The aim of this study was to examine how temporal and spatial electrophysiological activity relates to subjective visibility, while minimizing the influence of report-related factors.

### Subjectively visible and invisible trials are distinguishable at a 50-ms SOA

First, we evaluated the effect of backward masking on stimulus visibility across different SOA conditions in the main task. The behavioral paradigm is shown in Fig. 1A and 1B, and the results are summarized in Fig. 1C and 1D. Participants were required to localize briefly presented stimuli (left or right) followed by a mask, and subsequently rated their perceptual experience using the PAS. At the shortest SOA (8 ms), stimuli were mostly categorized as generating “no experience” (PAS 1: 79.1%), while at the longest SOA (150 ms) participants consistently reported some stimulus visibility (PAS 2–4: 96.8%). There was no significant difference in PAS responses between the mask-only condition (no target presented) and the 8 ms SOA condition (paired sample t-test: t(90) = −1.31, p =.193, Cohen’s d = −0.14). Crucially, the 50 ms SOA condition gave a mix of perceived and non-perceived trials (PAS 2–4: 70.1%; PAS 1: 29.9%, Fig. 1C), enabling a clear separation between subjectively visible (blue) and subjectively invisible (red) trials, as visualized in Fig. 1D.

Localization performance, assessed using d’, varied significantly across experimental conditions (F(2.39, 214.90) = 690.77, *p* <.001, η² = .89, Fig. 1D), with accuracy increasing as SOA increased. Bonferroni-corrected post-hoc comparisons confirmed significant differences in localization d’ between all SOA conditions (all p_bonf_ <.001; Suppl. Table 1). Performance was near chance at the 8 ms SOA (d′ = 0.07), and highest at the 150 ms SOA, significantly exceeding all other conditions. (d′ = 3.22), aligning with subjective awareness reports. Within the 50 ms SOA, subjectively visible trials (PAS 2–4) showed significantly higher accuracy (d′ = 2.51) compared to subjectively invisible trials (d′ = 1.03; p < .001; Suppl. Table 2).

These findings demonstrate that physically identical stimuli can produce distinct accuracy levels across SOA conditions. Specifically, within the 50 ms SOA, behavioral performance varies with subjective awareness, with significantly higher accuracy when participants report some level of awareness (PAS 2–4). The analyses reported here focus on the subjective experimental conditions (subjectively visible vs. subjectively invisible).

### EEG decoding reveals early time window of subjective visibility independent of report

To isolate neural correlates of subjective visibility from post-perceptual processing effects, we used multivariate pattern analysis (MVPA). MVPA offers greater sensitivity over traditional univariate approaches by leveraging distributed patterns of neural activity and their covariance structure (Haynes & Rees, 2006; Sandberg et al., 2014).

As outlined in Methods (Table 1), main task k-folding involved training and testing classifiers to estimate stimulus location from EEG data collected during the main task (including report), thereby incorporating post-perceptual processing effects into decoding results. In contrast, for the cross-decoding analysis, classifiers were trained on data from the no-report task and tested on the main task, enabling decoding of stimulus location in both subjectively visible and invisible conditions while minimizing report-related confounds. Decoding accuracy for all four experimental conditions, including objective (reference) conditions is provided in the supplementary materials (Suppl. Fig. 2).

Stimulus location could be reliably decoded above chance in both subjectively visible and invisible trials using k-folding within the main task (Fig. 2A and Suppl. Table 3). In subjectively visible trials, two prominent peaks in decoding emerged: the first at approximately 150 ms and the second at 230 ms post-stimulus onset.

**Figure 2.**
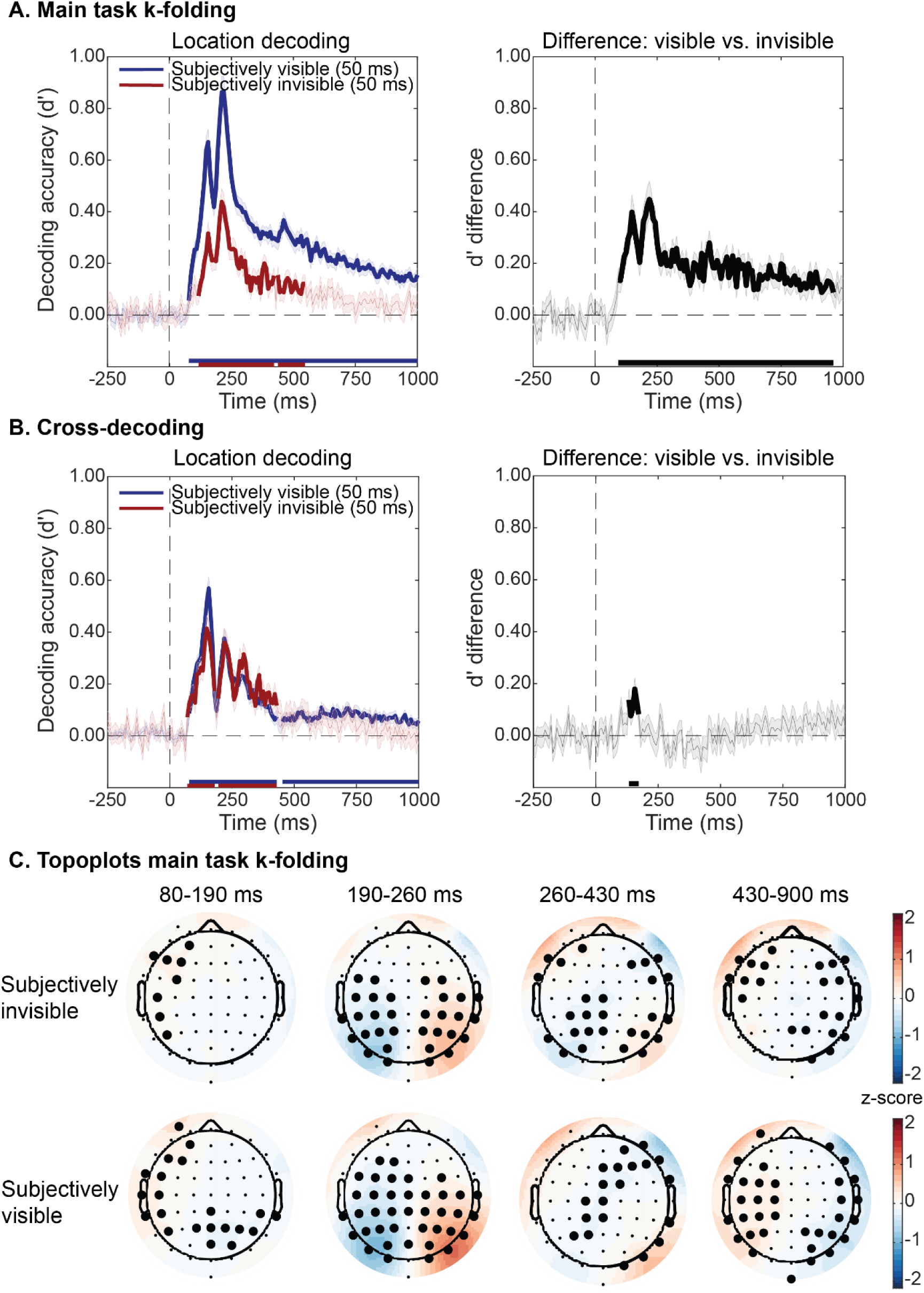
Location decoding (left vs. right) as a function of subjective visibility. **(A)** Main task k-folding (classifier trained and tested on main task data): group-averaged decoding performance (d′ ± SEM) for subjectively visible (blue) and invisible (red) trials (N = 83). Difference waves (visible minus invisible) are displayed next to each decoding plot. Thick lines indicate p < 0.05 (two-sided cluster-based permutation tests; against baseline for decoding, between conditions for differences). **(B)** Cross-decoding: classifier trained on no-report data and tested on main task data (N = 91), shown in the same format. **(C)** Haufe-transformed topographical maps at peak decoding intervals from the main task, showing standardized activation pattern coefficients. Thick electrodes mark significant clusters (p < 0.05). For statistical details, see Suppl. Table 3.

A direct comparison between subjectively visible and invisible trials revealed significantly higher decoding accuracy for subjectively visible trials, with differences emerging between 94 and 961 ms following stimulus onset. This contrast reflects differences in neural signals associated with subjective visibility, although it still includes report-related contributions.

Crucially, cross-decoding also showed decoding accuracy significantly above chance in both subjectively visible and invisible conditions (Fig. 2B and Suppl. Table 3), although with reduced overall accuracy. Importantly, the temporal window during which localization-related neural patterns could be decoded to distinguish subjectively visible from invisible trials (i.e., higher accuracy for visible than invisible conditions) was restricted to 133–172 ms (P < 0.05, Suppl. Table 3) post-stimulus onset. Decoding accuracy was significantly reduced at later processing stages (>175 ms) when the classifier was trained on no-report data. This window is substantially narrower than that observed in the main task k-folding analysis. This early window reflects a report-independent neural correlate of subjective visibility, reflecting enhanced neural encoding of consciously perceived compared to unperceived stimuli.

Topographical maps of classifier weight patterns during the main task k-folding further confirmed this result. Averaged over 80–190 ms, a window that includes the significant decoding period identified in the cross-decoding analyses, posterior electrodes in the right hemisphere contributed strongly in the subjectively visible but not the subjectively invisible condition (Fig. 2C). These patterns, derived via the Haufe et al. (2014) transformation of weight vectors, reflect univariate differences between stimulus classes. For reference, Supplementary Figure 3 shows maps for all main-task conditions and cross-decoding.

Together, these findings indicate an early, transient neural signature of subjective visibility (∼130–175 ms) that is independent of report-related activity. Later decoding effects observed in the main task k-folding are likely influenced by post-perceptual processes, highlighting that conscious perception in this paradigm is primarily associated with early posterior neural representations rather than sustained or frontal activity.

### Posterior, but not frontal brain regions, correlate with subjective visibility independent of report

Having established the temporal profile of report-independent subjective visibility, we next examined its spatial organization. While whole-scalp decoding captures distributed information, it does not indicate which regions contribute most strongly. We therefore performed region-specific cross-decoding using predefined electrode pools.

Classifiers were trained on the no-report task and tested on the main masking task, separately for occipital, temporal, parietal, central, and frontal electrode groups (Fig. 3). Training and testing focused on the 75–190 ms window identified by the cross-decoding analysis. Only diagonal decoding was used to isolate region-specific contributions.

**Figure 3.**
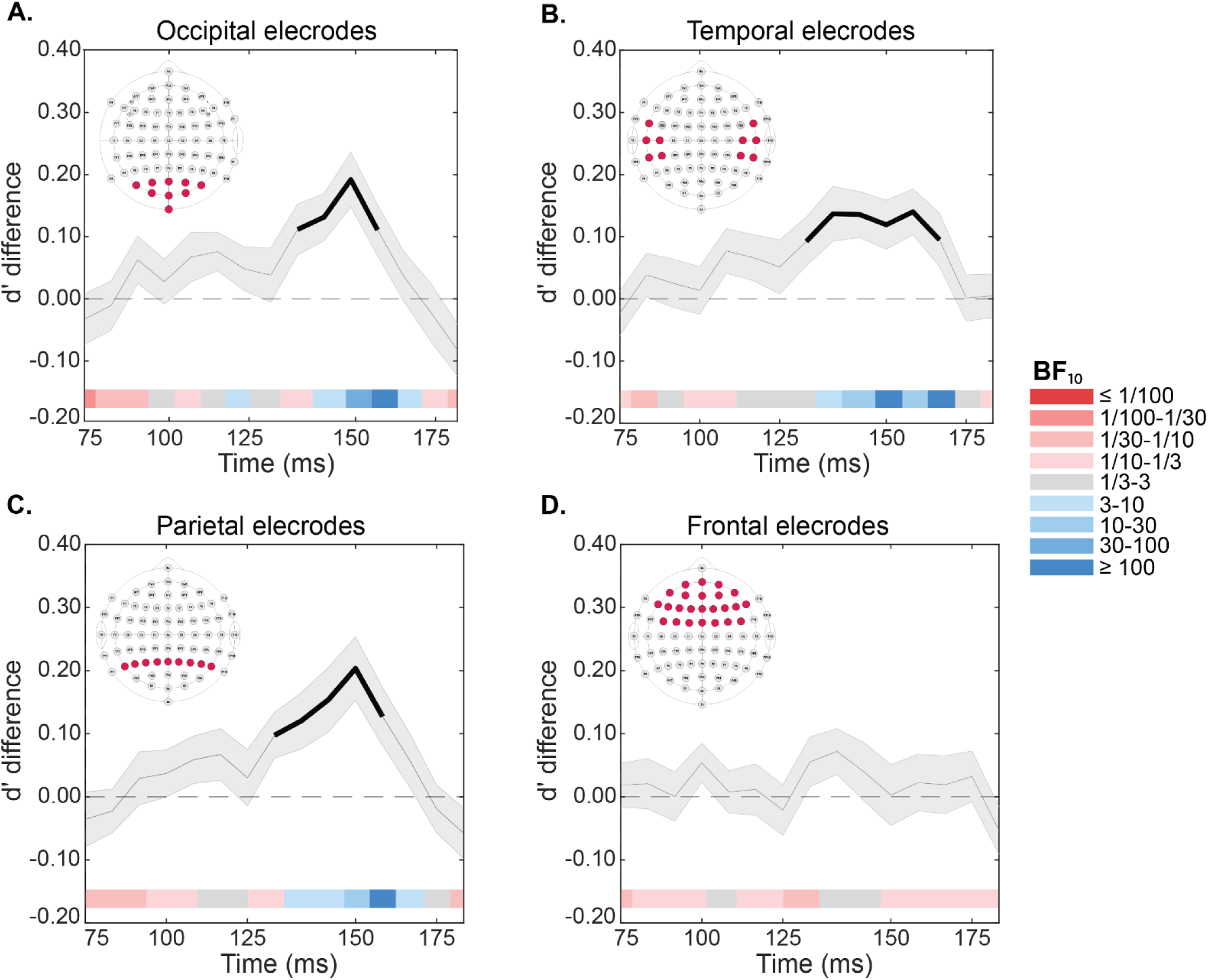
Spatial distribution of cross-decoding accuracy across cortical regions. Panels A–D show the difference in decoding d′ (subjectively visible minus subjectively invisible trials) for stimulus location (left vs. right) within the 75–190 ms time window for each channel pool: occipital (A), temporal (B), parietal (C), and frontal (D). Channel pool locations are indicated in the top-left corner of each panel. Classifiers were trained on no-report data and tested on main task data (cross-decoding; N = 91). Bayesian evidence is shown in the bottom bars (red/blue), illustrating the support for or against a difference in decoding between visibility conditions. Thick lines indicate p < 0.05 (two-sided cluster-based permutation test between conditions); shaded areas represent 95% confidence intervals. Significant decoding differences were observed in occipital, temporal, and parietal, but not frontal electrodes.

Significant decoding differences between subjectively visible and invisible trials were observed in the occipital (141–164 ms; Fig. 3A), temporal (133–172 ms; Fig. 3B), and parietal (133–164 ms; Fig. 3C) electrode pools, matching the early cross-decoding window. In contrast, classifiers trained on frontal or central electrodes showed no significant visibility-related differences (Fig. 3D).

Bayesian analyses confirmed region-specific effects of subjective visibility, as indicated by the colored bars in corresponding figures. In the early 130–175 ms window, occipital (median BF₁₀ = 7.17) and parietal (median BF₁₀ = 7.56) electrodes showed moderate-to-strong evidence for greater decoding accuracy in the visible compared to invisible condition, with temporal electrodes even stronger (median BF₁₀ = 17.97). Frontal electrodes (median BF₁₀ = 0.22) provided moderate evidence for the absence of an effect.

Overall, these findings demonstrate that early sensory regions primarily support visibility-dependent neural representations, while frontal areas do not reliably distinguish visible from invisible stimuli.

### Temporal generalization reveals early, transient correlates of subjective visibility

Previous research has linked conscious perception to metastable neural representations (Salti et al., 2012; Schurger et al., 2015). We therefore assessed whether stimulus representations associated with subjective visibility exhibit sustained temporal generalization.

Figure 4 shows the average temporal generalization matrices for subjectively visible and invisible trials, as well as their difference, for both the main task k-folding (including report) and cross-decoding (report-independent) analyses. Significant clusters were identified using one-sided permutation tests with a cluster-defining threshold of p < 0.01 and cluster-level correction at p < 0.01 (dark contours).

**Figure 4.**
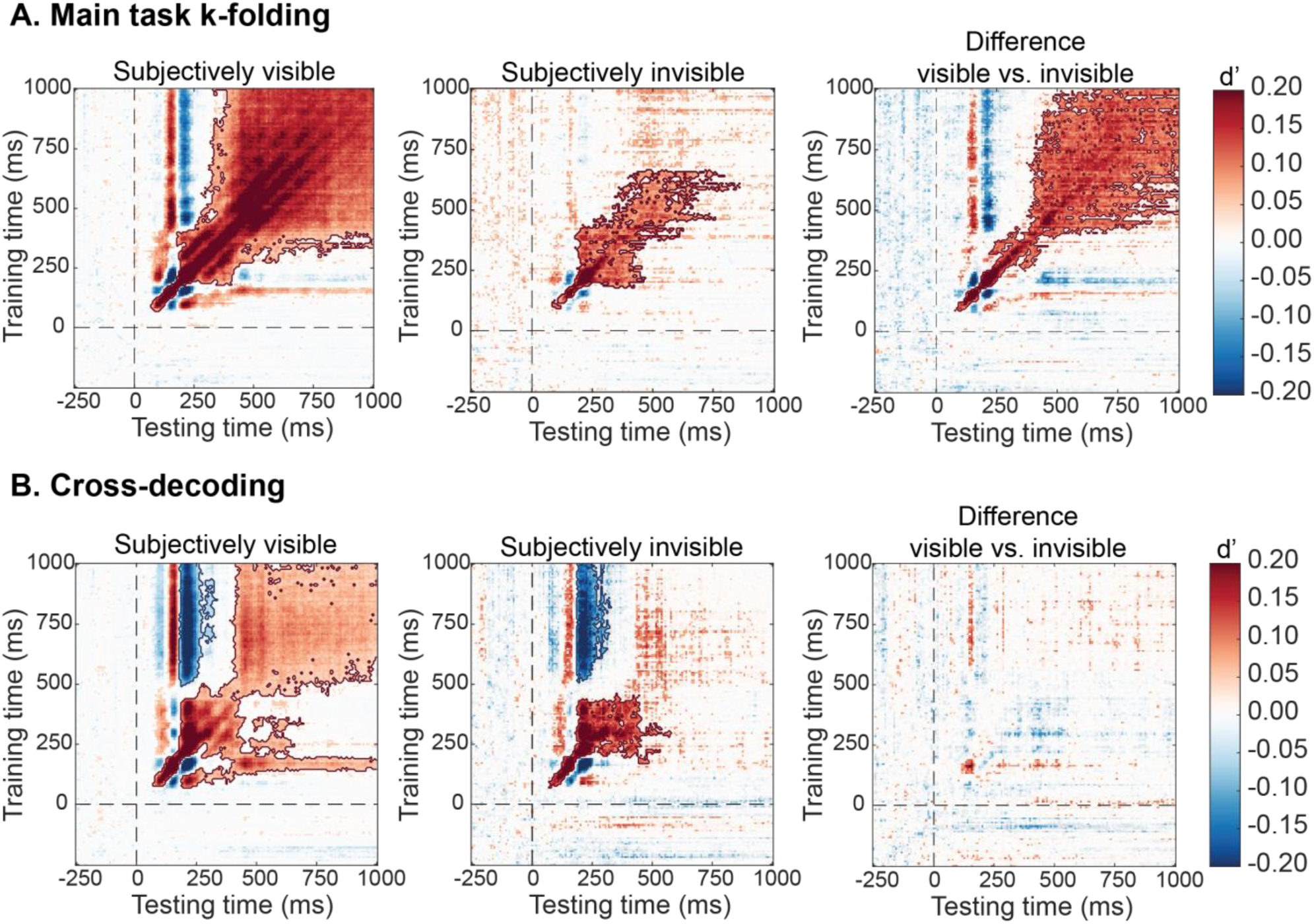
Temporal Generalization Matrices (TGMs) for decoding stimulus location. TGMs show classifier performance (d′) as a function of training time (y-axis) and testing time (x-axis), with color indicating decoding accuracy of stimulus location (left vs. right). **(A)** Main task k-folding: TGMs for subjectively visible, subjectively invisible trials, and their difference (visible minus invisible), respectively, when classifiers were trained and tested on main task data (*N* = 83). **(B)** Cross-decoding: TGMs when classifiers were trained on no-report data and tested on main task data (N = 91).

In the main task k-folding, subjectively visible trials showed a pronounced metastable pattern beginning around 200 ms and persisting from approximately 350 ms to the end of the trial (Fig. 3A), consistent with a stable neural representation. This sustained pattern was substantially stronger for visible than invisible trials. When subtracting invisible from visible trials, a prominent late difference emerged beginning around 100 ms (Fig. 4A, rightmost panel).

In contrast, cross-decoding revealed much weaker and shorter-lived temporal generalization (Fig. 4B). Subjectively visible trials showed limited sustained generalization between approximately 157–200 ms and again later in the trial, whereas invisible trials showed only brief generalization between 200–400 ms. Critically, when visible and invisible trials were directly contrasted using cross-decoding, the late metastable pattern disappeared (Fig. 4B, rightmost panel). Instead, significant decoding was restricted to the diagonal, indicating transient rather than sustained neural representations in a narrow window around 170–200 ms post-stimulus onset.

## Discussion

The goal of this study was to advance our understanding of the neural correlates of conscious perception by integrating report-based and report-independent paradigms within a multivariate cross-decoding framework. Using this approach, we identified a distinct early time window between approximately 130 and 170 ms after stimulus onset during which activity in posterior, but not frontal, EEG channels reliably distinguished subjectively visible from invisible stimuli. These results align with previous findings from no-report paradigms (e.g., Cohen et al., 2020; Hatamimajoumerd et al., 2022), demonstrating that while conscious content can still be decoded without report, classifier performance is markedly reduced, particularly in later time windows. Because this effect was obtained using classifiers trained on a no-report task and tested on a report-based masking task, it reflects neural representations that generalize across reporting contexts and are therefore minimally contaminated by post-perceptual processes like decision, motor, or introspective processes. These findings support theories that locate conscious contents in early recurrent processing within posterior cortex, such as Integrated Information Theory and Recurrent Processing Theory (Lamme, 2006; Tononi, 2008, 2012, 2012), and challenge accounts that locate the primary neural basis of conscious perception in prefrontal cortex (Dehaene & Changeux, 2011; Lau & Rosenthal, 2011; Mashour et al., 2020).

A central contribution of this study is methodological. By training classifiers on an independent no-report task and testing them on trials sorted by subjective visibility in a masking task, we were able to isolate content-specific neural representations that are both perceptually meaningful and independent of report. This hybrid strategy avoids a key limitation of standard report paradigms, in which neural signals related to perception are entangled with task relevance, decision formation, and response preparation (Aru et al., 2012; Fazekas & Overgaard, 2018; Tsuchiya et al., 2015; Naccache et al., 2015), while also overcoming a major weakness of pure no-report designs, namely the absence of a trial-by-trial measure of subjective experience. By using stimulus location as a content-specific proxy and PAS ratings only as test labels to assess how neural patterns relate to subjective visibility rather than as training signals, we provide a principled way to study neural correlates of subjective visibility that generalize across task contexts (report vs. no-report design).

In relation to previous work, our findings extend recent no-report and cross-decoding studies (Cohen et al., 2020, 2024; Hatamimajoumerd et al., 2022) by combining report-independent training with subjective trial labeling, allowing us to more precisely isolate visibility-related signals from confounds introduced by reporting and post-perceptual processed induced by task structure. Integrating subjective visibility ratings with cross-decoding of unreported stimuli thus represents a significant advance over existing no-report approaches.

Our results also speak to the interpretation of late centro-parietal ERP components, particularly the P3b, which has historically been associated with conscious access but is now widely thought to reflect post-perceptual processes (Aru et al., 2012; Pitts et al., 2014; Sergent et al., 2005). In line with recent no-report studies (Cohen et al., 2020; Hatamimajoumerd et al., 2022), we observed robust decoding accuracy in a later time window corresponding to the P3b in the report-based analyses, but a strong reduction in late decoding and metastable generalization when classifiers were trained on no-report data. This pattern confirms that the P3b and related late ERP components primarily reflect post-perceptual and report-related processes rather than perceptual visibility itself.

At the same time, our findings do not decisively falsify Global Neuronal Workspace Theory (GNWT) or fully validate posterior-only theories. Scalp EEG has limited spatial resolution and may be insensitive to distributed or deep prefrontal activity that could still contribute to conscious perception, which would require fMRI or intracranial recordings to detect. Moreover, the early posterior signal observed here could correspond to an initial ignition phase that precedes later global broadcasting, as proposed by GNWT (Dehaene & Changeux, 2011; Mashour et al., 2020).

Furthermore, our results may be compatible with Sergent’s “global playground” proposal (Sergent et al., 2021), which posits a transient, flexible network enabling the temporary sharing and maintenance of sensory representations without specific need for an action. In this view, conscious access can occur within a limited subset of the broader global workspace architecture, sufficient to sustain subjective experience but not necessarily to support decision-making or task execution. When task demands arise, as they do in our main task but not in the no-report paradigm, this playground may be further recruited by executive and decisional mechanisms, thereby transitioning into a fully engaged global workspace.

Nevertheless, the absence of any frontal contribution to cross-decoded visibility effects places important constraints on interpretations of late fronto-parietal activity. Our results add to accumulating evidence that the P3b and related late components primarily reflect post-perceptual processes such as decision-making, task relevance, and report preparation rather than perceptual visibility itself (Aru et al., 2012; Cohen et al., 2020; Hatamimajoumerd et al., 2022; Pitts et al., 2014; Schelonka et al., 2017; Shafto & Pitts, 2015). The disappearance of late metastable patterns under report-independent decoding further supports this view, indicating that sustained late activity is influenced by reporting and task demands rather than on the presence of a conscious percept.

Several limitations of our study should be acknowledged. Because classifiers were trained on an unmasked no-report task and tested on masked report trials, they may underestimate the full neural signature of subjective visibility if the two tasks recruit partially different representations. Our focus on stimulus location also captures only a limited aspect of perceptual content, potentially biasing decoding toward early sensory representations. As with all no-report approaches, we cannot entirely exclude internal monitoring or decisional processes, the “bored monkey” problem (Block, 2019).

In conclusion, our findings indicate that subjective visibility is associated with an early posterior neural signal that generalizes across task design (report and no-report), emerging between 130 and 170 ms after stimulus onset, with no detectable contribution from frontal EEG channels. By combining subjective visibility ratings with report-independent cross-decoding, we provide a powerful framework for isolating perceptual correlates of conscious experience while minimizing post-perceptual confounds. This approach offers a promising path forward for resolving long-standing debates about where and when conscious perception arises in the brain.

## Data availability statement

All code and data related to the present paper are available at UvA Figshare (link to be added upon acceptance).

## Author contribution

T.S. and V.A. conceptualized and designed the study. V.A. was responsible for data collection. A.H., S.G., and V.A. conducted the data analyses with guidance and supervision from J.F., J.C.F., U.O., and T.S. S.G. prepared the manuscript. All authors contributed to the revision of the manuscript and approved the final version.

## Declaration of interests

The authors declare no competing interests.

## Supporting information

Supplementary Figure 1. Location decoding accuracy (AUC) for subjectively visible vs. invisible trials.

Supplementary Figure 3. Scalp topographies showing standardized activation pattern coefficients

## Acknowledgments

We gratefully acknowledge Johannes Fahrenfort for his invaluable comments on the manuscript, guidance, and assistance with the ADAM toolbox. We also thank Lucija Blazevski and Carlotta Koch at the University of Amsterdam for their dedicated efforts in collecting part of the EEG data.

## Funding Information

This project was made possible through the support of a project grant from Amsterdam Brain and Cognition to UO and TS, and of a grant from the Templeton World Charity Foundation, Inc (TWCF-2022-30261) to UO and TS. The opinions expressed in this publication are those of the authors and do not necessarily reflect the views of Templeton World Charity Foundation, Inc. https://www.templetonworldcharity.org/. The funders had no role in study design, data collection and analysis, decision to publish, or preparation of the manuscript.

## Supplementary materials

### Behavior

**Supplementary Table 1.** Post hoc comparisons examining the effect of different SOAs on localization accuracy (*d′*).

| | | Mean difference ($d'$) | SE | t | P<sub>bonf</sub> |
| --- | --- | --- | --- | --- | --- |
| **8 ms** | *50 ms Invisible* | -0.967 | 0.077 | -12.607 | <.001 |
|  | *50 ms Visible* | -2.448 | 0.077 | -31.916 | <.001 |
|  | *150 ms* | -3.157 | 0.077 | -41.156 | <.001 |
| **50 ms Invisible** | *50 ms Visible* | -1.481 | 0.077 | -19.309 | <.001 |
|  | *150 ms* | -2.190 | 0.077 | -28.549 | <.001 |
| **50 ms Visible** | *150 ms* | -0.709 | 0.077 | -9.240 | <.001 |
| *Note.* P-value adjusted for comparing a family of 6 |  |  |  |  |  |

**Supplementary Table 2.**
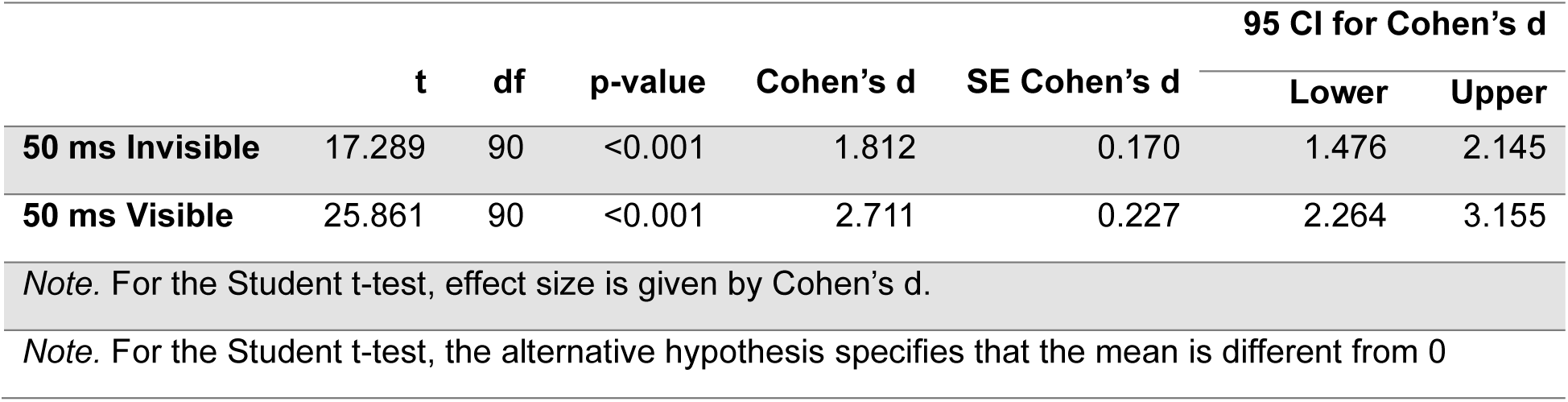
One-sample *t*-tests comparing localization *d′* to chance level for 50-ms SOA conditions (subjectively visible vs. invisible).

|  | t | df | p-value | Cohen's d | SE Cohen's d | 95 CI for Cohen's d |  |
| --- | --- | --- | --- | --- | --- | --- | --- |
|  |  |  |  |  |  | Lower | Upper |
| **50 ms Invisible** | 17.289 | 90 | <0.001 | 1.812 | 0.170 | 1.476 | 2.145 |
| **50 ms Visible** | 25.861 | 90 | <0.001 | 2.711 | 0.227 | 2.264 | 3.155 |
| *Note.* For the Student $t$-test, effect size is given by Cohen's $d$. | | | | | | | |
| *Note.* For the Student $t$-test, the alternative hypothesis specifies that the mean is different from 0 | | | | | | | |

### MVPA: EEG decoding does not reveal significant differences in subjective awareness independent of report when using AUC instead of d’

**Supplementary Figure 1.**
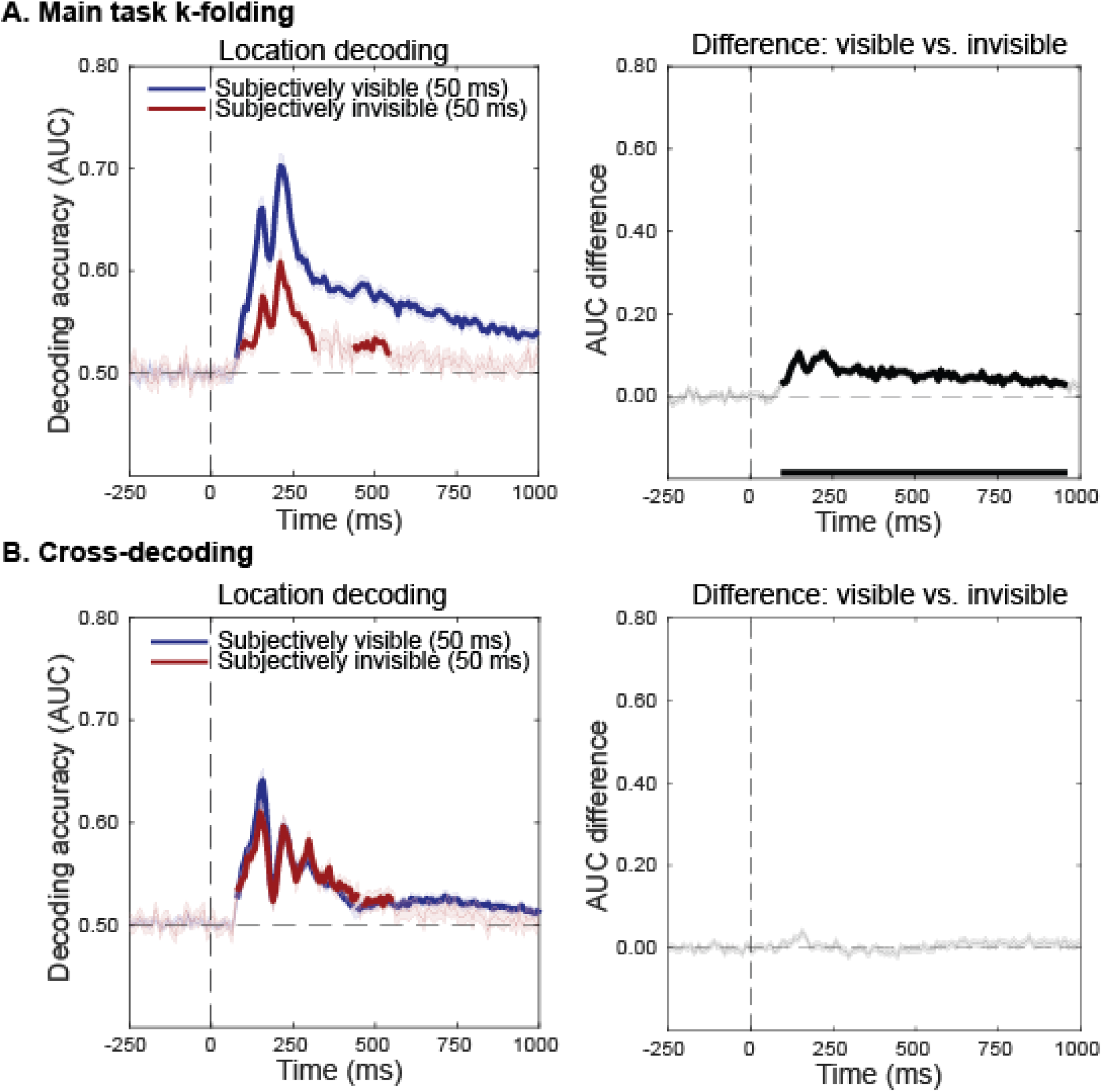
Location decoding accuracy (AUC) for subjectively visible vs. invisible trials. **(A)** Main task k-folding: Group-averaged diagonal AUC ± SEM for visible (blue) and invisible (red) trials using main task k-folding. Rightmost panel: difference between visible and invisible trials (visible − invisible). **(B)** Cross-decoding: Group-averaged diagonal AUC ± SEM for cross-decoding (classifier trained on localizer, tested on main task. Rightmost panel: difference between visible and invisible trials. *Thick lines indicate time points with p < 0.05 (two-sided cluster-based permutation tests; against baseline in A, C; between conditions in B, D)*

**Supplementary Figure 2.**
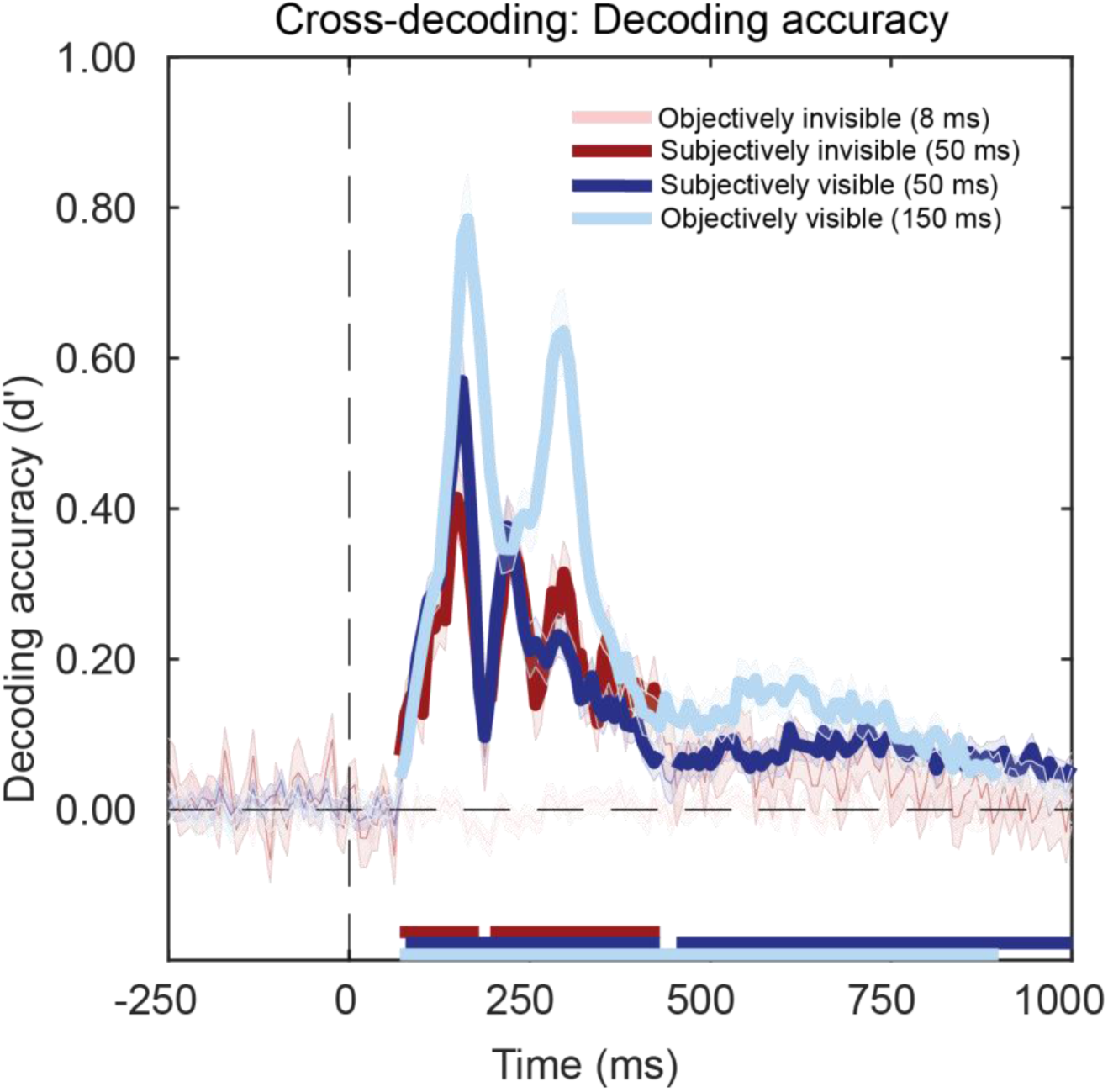
Group-averaged decoding accuracy (d′ ± SEM) across all conditions: subjectively visible, subjectively invisible, objectively visible, and objectively invisible. Cross-decoding was performed with classifiers trained on the no-report task and tested on main task data (N = 91). *Thick lines indicate time points with p < 0.05 (two-sided cluster-based permutation tests).*

**Supplementary Table 3.**
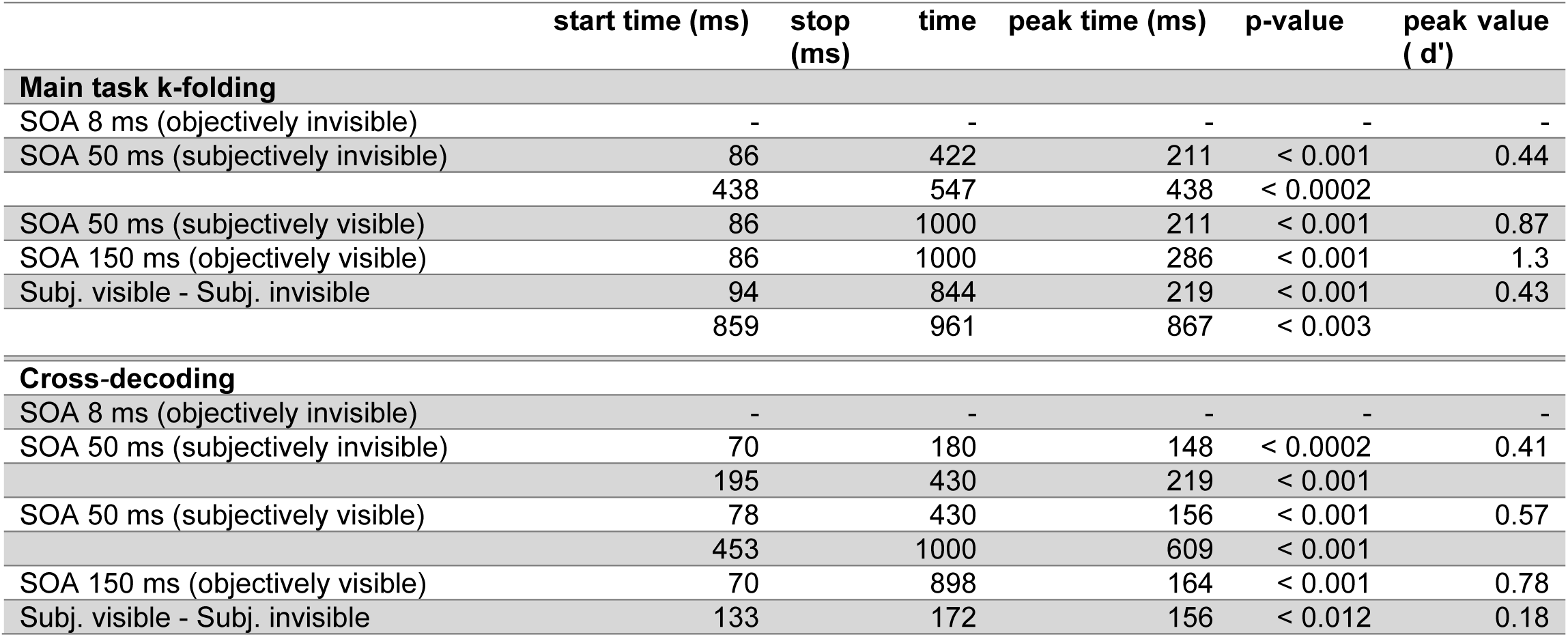
Significant results from diagonal decoding analyses.

**Supplementary Figure 3.**
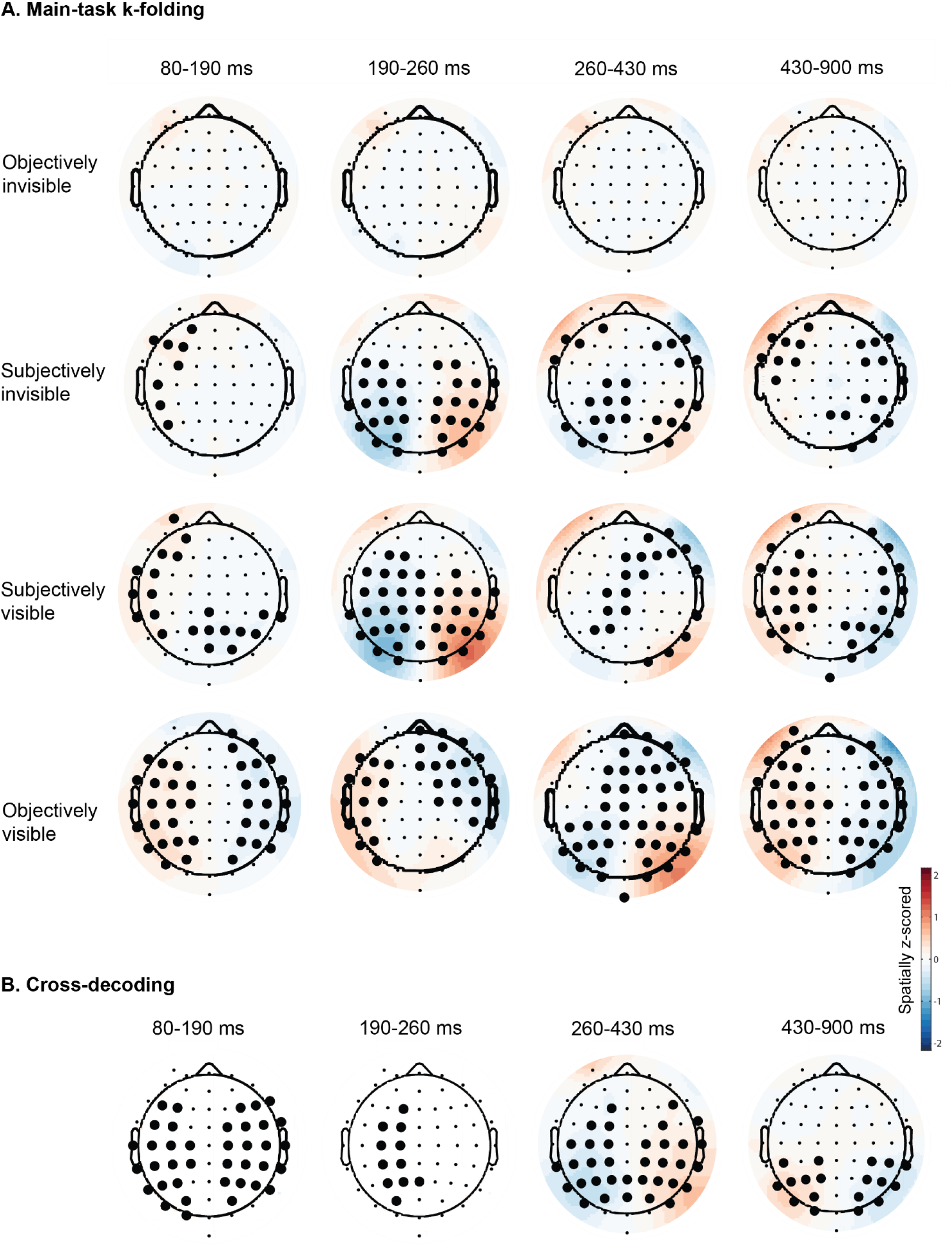
Scalp topographies showing standardized activation pattern coefficients. **(A)** Main task k-folding decoding for all conditions. Classifier trained and tested on main task data with PAS ratings. **(B)** Cross-decoding across time windows (80–190 ms, 190–260 ms, 260–430 ms, 430–900 ms), combining all conditions; classifier trained on no-report task without PAS ratings.

## Notes

### Competing Interest Statement

The authors have declared no competing interest.

