## Supplementary figures and images for "Posterior but not frontal neural signatures of subjective visibility in report-independent EEG decoding"

### Supplemental Data 1

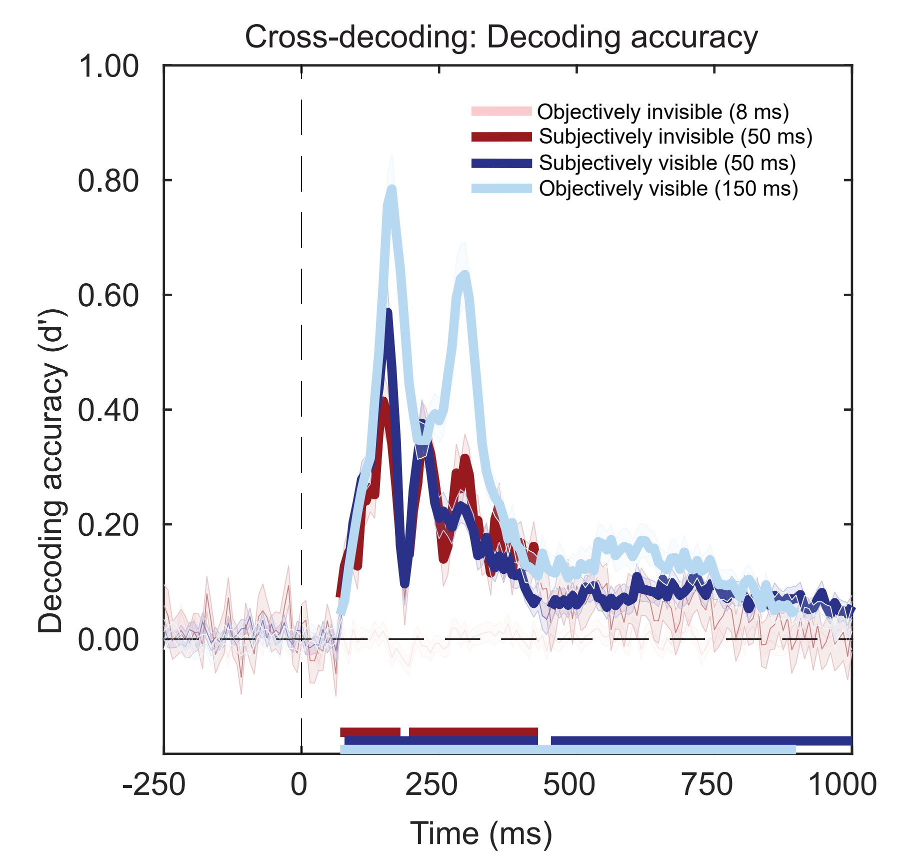

### Supplementary Figure 1. Location decoding accuracy (AUC) for subjectively visible vs. invisible trials.

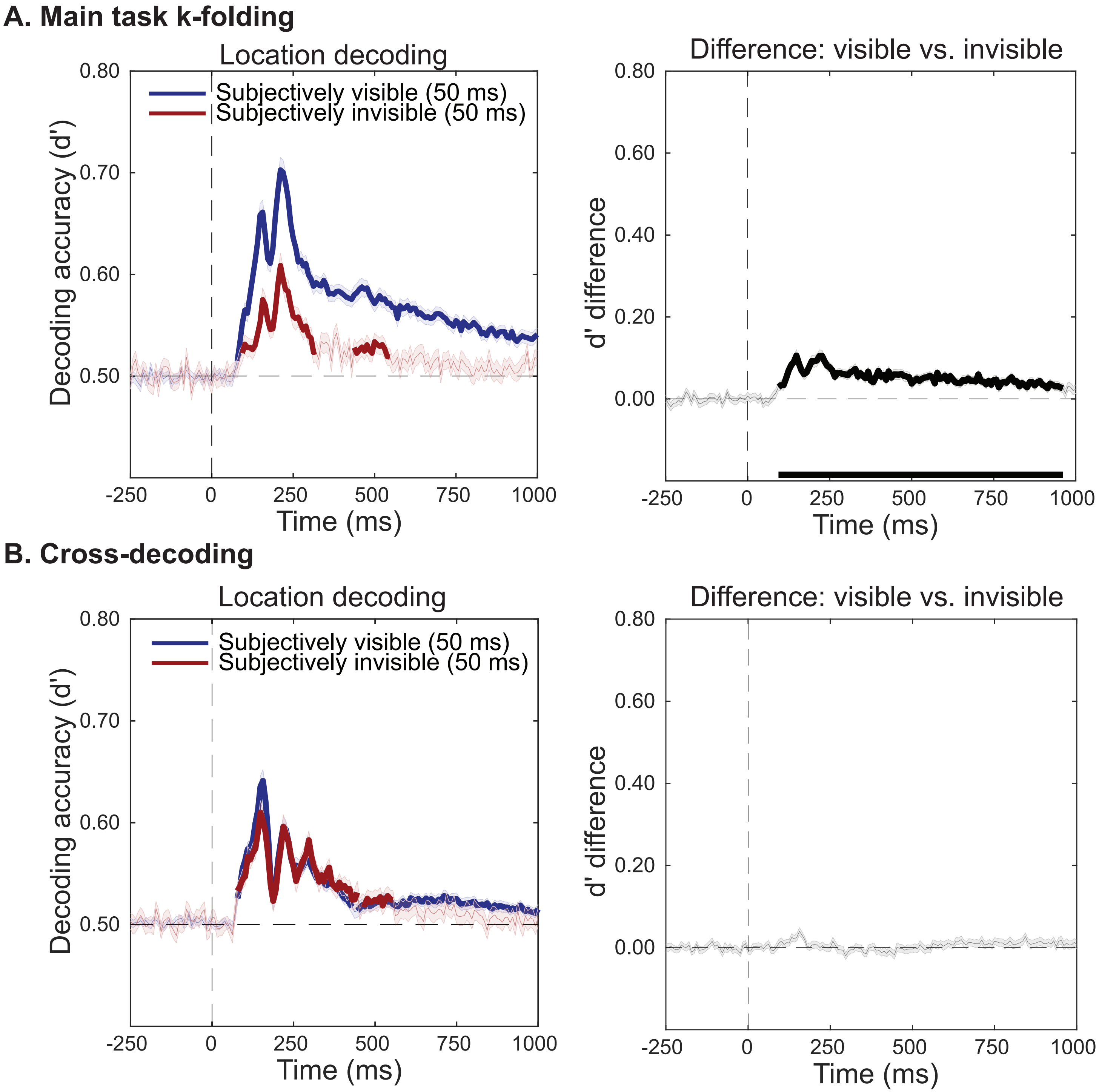

### Supplementary Figure 3. Scalp topographies showing standardized activation pattern coefficients

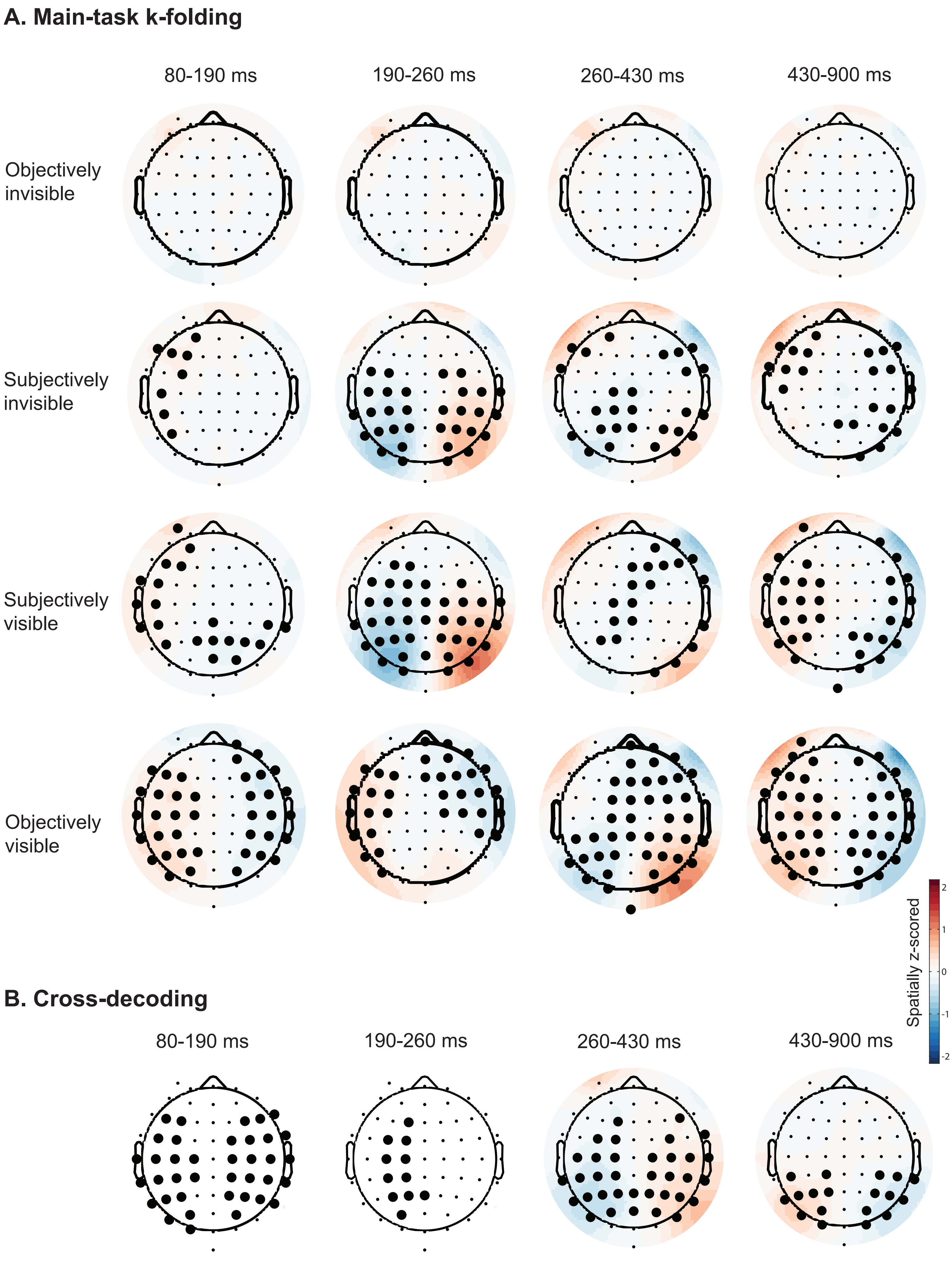
